# *Partners in health?* Investigating social genetic effects for married and cohabiting couples

**DOI:** 10.1101/688523

**Authors:** Jornt Mandemakers, Kasper Otten

## Abstract

‘Social contagion’ research suggests that health behaviors (BMI, smoking, drinking, etc.) spread through social networks, including dyadic ties such as between married/cohabiting partners. However, separating contagion from assortative mating (‘like seeks like’) and shared environmental factors remains notoriously difficult in observational studies. It is not possible to obtain exogenous variation in long-term partnerships (‘random mating’), but genetic approaches can offer a novel way to examine partner similarity and the role of social contagion. This paper explores possible social genetic effects among partners, i.e., effects of the partner’s genes on one’s own behavior. We use the longitudinal Health and Retirement Study with data on health behavior and genomic data for both ego and his/her partner to examine social genetic effects for BMI, drinking, and smoking behavior. For each outcome, we find support for social genetic effects. Americans of European descent were more overweight if they had partners with higher polygenic scores for BMI net of their own polygenic score. Similar findings were found for the number of drinks per week and cigarettes per day. Longitudinal analyses that conditioned on past health behavior of both spouses confirmed these findings. We further explored whether susceptibility to the partner’s influence differed between men and women, but did not find consistent differences across outcomes. Findings are further discussed in the light of ramifications of social genetic effects for the social and biological sciences.

---

There is a growing interest in the possible consequences of the ‘social genome’; the impact of genotypes of others in our direct environment on our own phenotypes. These effects have been referred to as social genetic effects (SGE) or indirect genetic effects^1^. Although SGE have been established for numerous types of animals^1-3^, trees^4^, and bacteria^5^, and are sometimes found to be even stronger than direct genetic effects^1^, they were until recently virtually ignored in studies on human genetics (but see these recent exceptions^6-8^). SGE have been reported for friends and schoolmates in US high schools for educational attainment and BMI (Body Mass Index)^6^, but SGE have not been studied for probably the most important relationship in adulthood; the partner. Despite the importance of the partner for our health and well-being and decades of research, our understanding of its role is still limited^9, 10^. This study provides a first attempt to examine the influence of the partners’ genome for one’s healthy lifestyle among married/cohabiting couples.

Previous research shows that the partner’s lifestyle has considerable associations with one’s own lifestyle across a variety of domains, including obesity, physical activity, eating, alcohol consumption, weight loss, and smoking^10, 11^. Partners share resources and they may exert a positive influence by promoting a healthy lifestyle^12, 13^ and by sanctioning unhealthy behavior^13-17^, although partners may also reinforce unhealthy behaviors. These findings align with the ‘social contagion’ strand of research, which shows that health behaviors spread through social networks, including dyadic ties such as between partners^12, 18, 19^. However, it is notoriously difficult even in longitudinal research designs to establish causality because of assortative mating (‘like seeks like’) and contextual confounding^20^. Although natural experiments have been used to examine social contagion effects (e.g., student dorms^21, 22^), they are rare, and we know of no such experiment for partners (fortunately by the way).

In this paper we offer a novel approach to study social contagion for health behaviors in long-term partnerships by exploiting data on the genetic profiles of both partners in married/cohabiting couples. Genetics offers a unique angle to examine the partner’s influence because genes are randomly assigned at conception given parental genotypes, are not directly visible, can be measured, and have an ongoing influence on the phenotype. This means that the partner’s genotype, as opposed to his/her phenotype, is immune to influences from the (couple) environment and reverse causality through influences from ego. Because married/cohabiting people are exposed to their partner’s behavior (ego’s environment), which is partly driven by the partner’s genes, we may be able to detect SGE. For instance, having a partner who is genetically predisposed to alcohol dependence may increase one’s own alcohol consumption because the partner ensures there is always alcohol available at home. What is more, SGE may be particularly likely for partners compared to other social ties because partners face large costs to leaving their relationship and are typically exposed to each other on a daily basis, which may increase incentives to influence the partners behavior and also the willingness to conform. Although genetics are implicated in partner selection^23-30^, by using data on both partner’s genetic make-up we can examine genetic assortative mating *and* control for it.

We use data from the Health and Retirement Study (HRS), a longitudinal study that follows a nationally representative sample of adults aged 50 and older in the United States every two years since 1992. DNA samples have been collected between 2006-2010 for over twelve thousand participants. If a participant has a partner, this partner is automatically selected to participate in the study as well, even if he or she is younger than 50 years. We examine three outcomes for which extensive information on genetics is available and that are important indicators of a healthy lifestyle; namely BMI^31^, as a measure of adiposity, and two measures of health behavior: the level of alcohol consumption (drinks per week, DPW)^32^, and smoking (cigarettes per day, CPD)^32^. We start by examining partner similarity on a phenotypic and genotypic level for these outcomes. Recent findings show weak genotypic similarity between partners for education, height, and BMI^23-30^, but to our knowledge no research examined drinking and smoking behavior.

We then present the two main analyses. First, are there are social genetic effects of the partner net of one’s own genetic effects? Measured genetic differences between partners offer a way to separate genetic influences of ego from alter (the partner). This analysis comes with the caveat that these SGE estimates cannot be strictly interpreted as causal because controlling for ego’s PGS will only partly control for selection. There may be other genetic and non-genetic factors that come into play with partner selection^28, 33^. These selection processes could indirectly lead to an association between the partner’s PGS and ego’s outcome, i.e., spurious SGE. In the second main analyses we therefore exploit the longitudinal nature of the HRS to control for selection by conditioning on initial observed levels of health behavior of ego and the partner. We further investigate the role of assortative mating by including controls for family background, height and socio-economic attainment. Because men and women differ in healthy lifestyle and genetic influences on health may be different for men and women^34^, we report pooled and sex-stratified analyses. In addition, recent findings show the importance of environmental factors in moderating the effects of one’s genetic predisposition for health outcomes (e.g., BMI^35, 36^). In this study the partner is the environment of interest, we therefore also investigate whether effects of one’s genes partially depends on the partner’s genes. A person’s genetic propensity to drink might for example be more influential when his/her partner has a higher genetic propensity to drink. Such interactive effects, have also been referred to as social epistatic effects (SEE)^6^.

## RESULTS

### How similar are partners?

We find that there is considerable similarity at the phenotypic level in health behavior but there is a much lower similarity at the genetic level. Correlations for observed health behaviors between married/cohabiting partners in the HRS were modest to large (BMI *r =* 0.23, DPW *r =* 0.48, and CPD *r =* 0.37). To examine genetic similarity we constructed a PGS for each of these three health behaviors based on the most recent GWAS findings^31, 32^ (see methods for details). We confirmed that the PGS were associated with their respective phenotypes. Standardized estimates of ego’s PGS on respective outcomes ranged from 0.334 (SE=0.013) for BMI to 0.210 (SE=0.012) for DPW and were lowest for CPD with 0.079 (SE=0.012) (also see Fig. 1). For BMI the genetic correlation between partners was similar to a previous study (*r* = .04)^24^. To our knowledge, previous research did not examine partner genetic similarity for drinking and smoking behavior. We find genetic correlations between partners for DPW and CPD to be slightly larger than for BMI (*r* = 0.05 for both, see further Fig. S1). The lower level of genetic compared to phenotypic similarity is to be expected given prior research^24-29^; (1) genes are fixed at conception, and (2) not directly visible, which makes genetic assortative mating less likely. Nevertheless, the relative absence of genotypic similarity is somewhat surprising, since selection based on heritable phenotypic traits could also indirectly lead to genotypic similarity^27^. The absence may be an indication that the phenotypic correlation in health behaviors between partners is driven more by contagion than selection.

**Figure 1.**
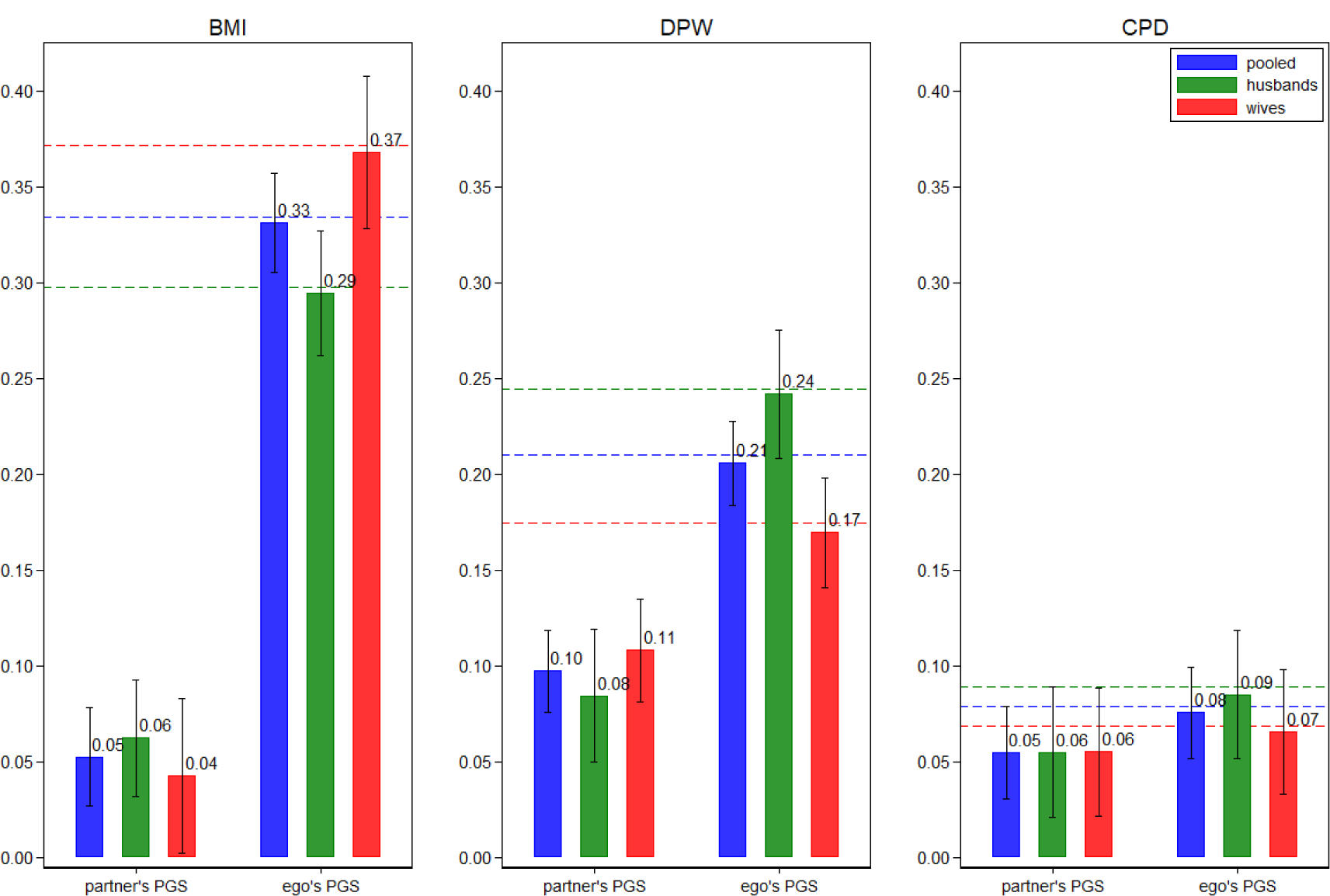
Social genetic effects of the partner. Estimates of social genetic effects of the partner (rounded to two decimals). Effect of partner PGS net of one’s own (ego’s) PGS for BMI, DPW, and CPD on associated outcomes (time-varying) in a pooled model and stratified by sex with socio-demographic controls and PC’s of ego and the partner. Outcomes are standardized as are the PGS. The figures also show the simultaneous effects of ego’s PGS in these models and the dashed lines (blue = pooled, green = husbands, red = wives) depict the baseline effect of ego’s PGS on the outcome in a base model with socio-demographic controls and PC’s of ego. CIs (95%) are robust to clustering within individuals and households.

### Is the partner’s genome associated with health behavior?

Estimates of social genetic effects of the partner in the HRS are summarized in Fig. 1 (detailed estimates are reported in Table S2-S3 in the SI appendix). We report the effects of the partner PGS net of ego’s PGS to control for genetic assortative mating in a pooled model to maximize statistical power and also separately by sex (referred to as husbands and wives for readability) to examine the possibility that (social) genetic effects differ by sex. The left figure shows the estimates for BMI, the middle for DPW and the right for CPD. All models include socio-demographic controls and principal components (PC) to correct for population stratification (see methods). We standardized the outcomes and the PGS to facilitate comparison. For each of the three outcomes we observe sizeable social genetic effects of the partner on ego’s behavior. For BMI an increase in the partner’s PGS with one standard deviation is associated with a 0.053 (SE=0.013) standard deviation increase in BMI (see Table S2), which is almost one sixth of the direct effect of one’s own PGS on BMI (0.331). For the number of drinks per week (DPW) the social genetic effect is about twice as large (0.097 (SE=0.011)) and almost one half of the direct effect of one’s own genes (0.206). The social genetic effect of the partner’s cigarettes per day (CPD) is about the same size as for BMI (0.055 (SE=0.012)) but this time the social genetic effects are even more than half the size of the direct genetic effects (0.076). Also note that the direct genetic effects of ego were only slightly reduced by including the partner’s PGS (compare dashed lines to ego’s PGS in Fig. 1), which suggests that direct and social genetic effects have independent effects.

It is often claimed that especially men’s health benefits from being in a long-term relationship^37^ because men may be more susceptible to social control of their partners and thus more likely to abstain from unhealthy behaviors. Stratifying the analysis by husbands (green) and wives (red), however, we see that the social genetic effects are fairly similar and the differences are not statistically significant. The direct genetic effects do differ, as the effect of one’s own PGS on BMI is stronger for wives. For DPW we observe the reverse pattern. While the direct genetic effect is lower for wives than husbands for CPD, the social genetic effect is roughly similar for husbands and wives.

Next we turn to social epistatic effects (SEE), i.e., interactive effects between one’s own genes and the partner’s genes. Positive SEE indicate that the partner’s genes reinforce the effect of one’s own genes because the partner’s genome may partly shape a more favorable environment to live out one’s genetic predisposition. For instance, the partners has a high PGS for CPD and so may be less likely to object to smoking at home. In theory, there could be negative SEE as well, but we think positive SEE is more likely for BMI/DPW/CPD because they have a large social component. We examined SEE directly by including interactions with ego’s and the partner’s PGS. Only for BMI we find statistically significant positive SEE; indicating that partners may reinforce each other’s genetic predispositions for BMI (see Figure S2 in the SI).

To control for selection not related to ego’s PGS, we carried out additional analyses that adjust for well-known patterns in assortative mating. We include educational level and interactions between education and sex, and mean parental education for both partners. There is ample evidence that education plays an important role in partner selection^27^, and there are large educational gradients in health behaviors^38^. We further include controls for height of both partners and interactions with sex, because it is also well known that height plays a role in partner selection^28^. We furthermore control for region of birth of both partners to take geographic proximity into account. Including this large set of controls reduces the estimates of the direct genetic effects (ego’s PGS) and the social genetics effects (partner’s PGS), but does not lead to substantially different results (see SI Table S2-S3).

### Is the partner’s genome associated with health behavior conditioning on past behavior?

As a second strategy to control for selection into partnerships we reran the analyses including both ego’s and the partners’ first observed scores for a phenotype (BMI/DPW/CPD respectively), thereby directly controlling for observed phenotypic selection. This approach should be seen as a conservative test of our hypothesis of social genetic effects. By conditioning on previous behavior we are effectively examining inter-individual *change* in health behavior in the period of observation, which severely reduces the amount of variation. And by using the HRS sample we examine change for a relatively old sample who are in long lasting relationships (aged ∼65 and in relationships of ∼34 years on average). There may be little change in health behavior in the period of observation given that most health behavior habits are formed at a much earlier age^39^. Perhaps the influence of the partner on health behavior has already largely played out before the period of observation.

Results of these conditional analyses are depicted in Figure 2 (see Table S3-S4 in the SI for details). By including both partners’ initial behavior, we first see much reduced effects of PGS of ego and the partner compared to the unconditional models (take note of the different scaling of the y-axis). As mentioned, these reduced estimates have a different interpretation, as now we are examining health behavior conditional on relatively recent behavior (between 2 and at most 14 years ago). The result of this more conservative test confirms the presence of positive social genetics effects for all three health behaviors. The pooled estimates for BMI (0.012(0.006), p=0.059), DPW (0.021(0.007), p=0.002), and CPD (0.013(0.007), p=0.059) are positive, although not significant in two-tailed tests for BMI/CPD. The pooled estimates for BMI/CPD are driven downwards by the lack of social genetic effects for wives as becomes clear by examining the sex-stratified estimates. For wives, the partner’s PGS estimates hover around 0 and are no longer significant. For husbands, we observe a decrease compared to Fig 1., but the estimates for BMI/CPD remain significant. Note that for these outcomes the social genetic effects of the partner are on par with that of the husband’s own direct genetic effects or even larger. For DPW, contrary to BMI/CPD, the social genetic effects are no longer significant for husbands, but they are for wives.

**Figure 2.**
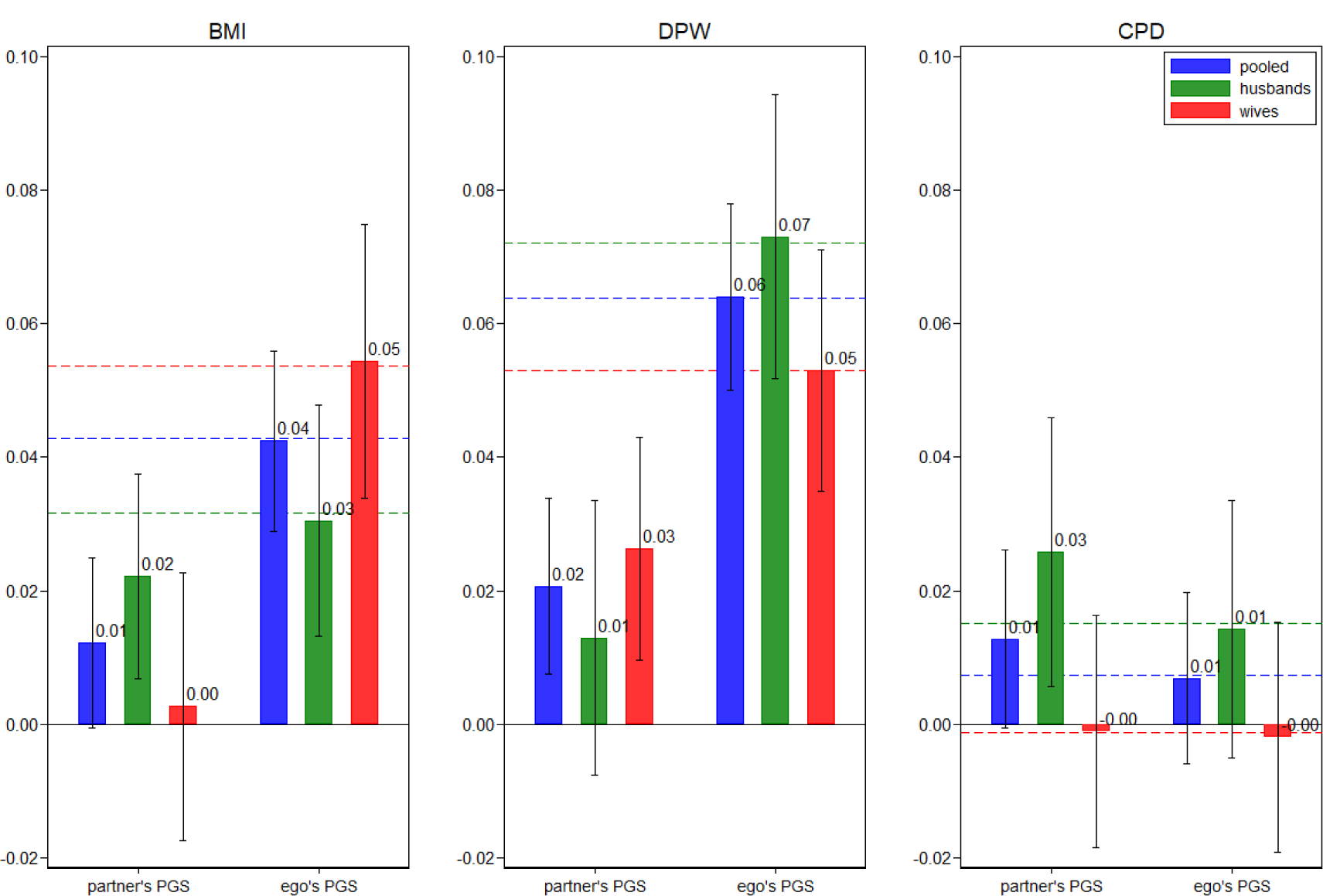
Social genetic effects of the partner conditioning on initial behavior of ego and partner. Estimates of social genetic effects of the partner conditioning on initial behavior of ego and partner (rounded to two decimals). Effect of partner PGS net of one’s own (ego’s) PGS and net of one’s own and the partner’s initial level of each outcome for BMI, DPW, and CPD on associated outcomes (time-varying) in a pooled model and stratified by sex with socio-demographic controls and PC’s of ego and the partner. Outcomes are standardized as are the PGS. The figures also show the simultaneous effects of ego’s PGS in these models and the dashed lines (blue = pooled, green = husbands, red = wives) depict the baseline effect of ego’s PGS on the outcome in a base model with socio-demographic controls and PC’s of ego and initial levels of the outcome for both ego and the partner. CIs (95%) are robust to clustering within individuals and households.

We examined social epistatic effects for these conditional models as well, but we do not observe SEE for BMI as we did before. In the conditional models we only observe SEE for CPD: the effect of ego’s PGS on CPD is stronger for people who have partners with a higher PGS for CPD (Figure S3 in the SI). Note that the effect is driven by husbands as the effect is only significant for men in sex-stratified analyses.

Finally, as may be expected, including the set of assortative mating controls makes a smaller difference to the conditional analyses than to the unconditional analyses (Table S4-S5), which confirms the idea that controlling for ego’s and the partner’s past behavior serves as an effective way to control for selection.

## Discussion

We examined social genetic effects of the partner for three widely studied health behaviors (BMI, drinking and smoking) using genome-wide data of both partners in married/cohabiting couples of the Health and Retirement Study, a large nationally representative sample of older American couples. We confirm previous research showing the existence of limited genetic similarity between partners for BMI^24^ and we provide novel estimates of partner genetic similarity for drinking and smoking behavior. For these last two health behaviors, we also find low levels of genetic similarity. There seems to be little genetic assortative mating when it comes to health behavior. We do observe robust social genetic effects (SGE), which offers an alternative genetically rooted explanation for the large phenotypic partner similarity in healthy lifestyle. People are more overweight, drink more, and smoke more, if they have partners with higher polygenic scores for these behaviors. In absolute terms the SGE are largest for drinking behavior. Our results suggest an important role for the partner’s genome. We find relatively large social genetic effects of the partner vis-a-vis the direct genetic influence of ego’s own genes.

Social genetic effects form a novel way to study the partner’s influence and the couple environment for health. This approach improves upon the reflection problem and reverse causality that hampers studies of social influence because the partner’s genotype, as opposed to his/her phenotype, is immune to influences from the (couple) environment and direct influences of ego. Our design controls for partner selection as alternative explanations for the presence of social genetic effects. We presented SGE net of direct genetic effects of ego, which controls for selection on genetic predispositions related to our outcome variables. Still SGE could be spurious because of selection on some other characteristic. We employed a broad set of socio-demographic controls for assortative mating and the SGE were only slightly reduced. Finally, most importantly, analyses that exploit the longitudinal nature of the HRS by conditioning on earlier health behavior of ego and the partner show that SGE largely remained.

Our paper revealed genetic cornerstones of social contagion within couples, but has several limitations that we hope future research will confront. First, the analyses were based on the HRS, which is a sample of older Americans who may be set in their ways and hard to influence in their behavior. Note that the couples were together for ∼34 years on average. The large difference between the SGE and the SGE conditional on previous health behavior of ego and partner may be seen in that light. The unconditional SGE may be relatively large given the large time for these effects to accrue, whereas the conditional findings may be conservative. We suspect that unconditional SGE will be smaller for younger populations and conditional SGE larger, but this remains an open question. Second, we used the most up-to-date GWAS to create polygenic scores for BMI, drinking and smoking^31, 32^. These captured about half of the SNP based heritability for the studied health behaviors, but still a large part is missing. PGS are likely to become more predictive as larger GWAS come out and so the potential to detect SGE will increase. We observed SGE for each health behavior, but we refrained from making statements about differences in the role of the partner for specific behaviors. It would be difficult to do so, as the size of the SGE also depends on heritability of an outcome and how well we can estimate the genetic influences and these differ among the three outcomes. Third, we limited the analyses to Americans of European descent because the polygenic scores were based on GWAS of people of European descent and are therefore less predictive in other ancestries. This choice limits the generalizability of our findings. Fourth, one’s PGS can be seen as an indirect measure of parental genotypes, so the SGE of the partner may also reflect possible SGE effects of parents-in-law (through socialization). We controlled for a set of family background controls, which reduces this possibility. However, maybe such effects should not be ruled out, because interpreting SGE as the effects of fixed predispositions of a partner on ego would include such socialization effects. Finally, we studied the three health behaviors in isolation and did not examine cross-trait effects. It is plausible that there may be cross-trait SGE given large phenotypic and genotypic correlations between BMI, drinking and smoking^32^ and the possibility that genes have ‘pleiotropic effects’ (having effects on multiple phenotypes)^40, 41^. A recent suggestion to study social influence effects using genes as instrumental variables forms another approach which may give insight here, but suffers from stricter assumptions^42^.

### Implications science

The presence of social genetic effects among partners deindividualizes the interpretation of genetic effects, which resonates with recent findings regarding SGE for educational attainment^6^. If a person’s phenotype is not just based on his or her own genes but also on the genes of peers, the assumption of independence of the units of analysis is violated. For example, the estimate of individual genetic effects could absorb the effect of partner’s genotype if the analyses do not account for partner’s genotype. This coupled with assortative mating could lead to substantial inflation of individual genetic effects. Although, our findings show that direct genetic effects were only slightly attenuated after including the partner’s PGS. More broadly, evidence for human social genetic and social epistatic effects would require rethinking current views of the meaning of polygenic scores in social scientific research. It also complicates the idea of gene-environment interactions taking place in a static non-reactive environment. Social epistatic interactions can be seen as gene-environment interactions, where the environment is made up of another individual whose behavior is partly influenced by his/her genes. In marriage/cohabitation it appears this the partner, but perhaps children and other household members should be considered as well. We found limited evidence of positive social epistatic effects, for BMI and in the conditional analyses for smoking. That does not rule out negative SEE, perhaps these are possible for competing outcomes; where one’s success diminishes returns for another. For instance, lazy people may thrive with thrifty partners and vice versa.

We focused on health behavior in married and cohabiting relationships. There is no reason why SGE would be limited to this tie only, as previous work already found SGE for schools and friends^6^. The environments (worksites, neighborhoods, schools, households, etc.) we navigate in life are populated by others who shape and mold these environments. These environments can be seen as genetic landscapes, which may give insights into the role colleagues^18^, neighbors^43^, and other household members^21, 44^. Moreover, besides healthy behaviors, other phenotypes could be susceptible to SGE, for instance, related to mental health and well-being^45^.

### Broader implications

The results contribute to a broader understanding of how health behaviors are molded and highlights the importance of people’s social surroundings. Why do some people keep smoking despite widespread knowledge of its harmful effects? The partner is likely important, but is that because of health selection which clusters health and wealth, because of shared environmental influences that affect both partners, or because partners reinforce each-other and keep unhealthy behavior locked in place? Our results suggest that this last reason is likely, and suggests an important role of the partner. Furthermore, this study has illustrated how genetic variation within and between couples can help to overcome difficulties in making causal inference regarding the role of the partner for health. Future applications could widen the evidential base of health prevention programs and/or medical care, as positive partner influence may lead to (unintended) positive spill-over effects on partners. These findings highlight the potential of targeting health interventions at couples, at partners of people who have already been identified as being at high risk, and at people most susceptible to social influence of the partner.

## Methods and Materials

### Data

The Health and Retirement Study (HRS) is a longitudinal household study that follows a nationally representative sample of adults aged 50 and older in the United States every two years since 1992^46^. We used the DbGap deposited DNA samples that were collected in 2006-2010. If a participant had a partner, this partner is automatically selected to participate in the study as well. We limited the sample to married/cohabiting couples of European decent where both partners had valid genomic data and excluded proxy interviews. Only couples of whom both partners were of European descent are included in the analysis because polygenic scores were based on European GWAS and have reduced predictive power in other ancestries. We further excluded observations of couples in case one or both partners was no longer living independently, but for instance in a nursing home. The analytical samples comprised ∼50,000 observations from waves 1992-2016 for ∼6,000 persons in ∼3,200 couples. Note that the sample differs somewhat between outcomes (see Table 1 in the SI for details). This study was conducted with institutional review board approval from Utrecht University, the Netherlands.

### Measures

All measurements are made for ego and partner. BMI (body mass index) is measured by dividing weight in kilograms (reported each wave) by length in meters squared (weight was reported each wave, height was taken from the first interview). Drinking is measured by the number of alcoholic drinks per week (DPW). Smoking was measured by the number of cigarettes per day (CPD) and non-smokers were set to zero. We take the natural logarithm for DPW and CPD, left anchored at 1 (ln(y+1)). All phenotypes were measured each wave, except for the first two waves that did not have comparable measures for DPW/CPD and could not be used for DPW/CPD.

Genetic propensity for BMI/DPW/CPD is measured by the use of polygenic scores (PGS). A PGS is the aggregation of many small genetic effects scattered across the genome on a phenotype. It is computed by weighting the alleles at the different loci across the genome with their association to the phenotype of interest, and then summing these weighted alleles. Information on the association between alleles and phenotypes is derived from recent large-scale publicly available GWAS^31, 32^ that did not overlap with the HRS. Following recommendations we calculated the PGS based on the genotyped SNPs of the 22 autosomal chromosomes, and do not account for linkage disequilibrium or p-value thresholding, as this was shown to have no or a negative effect on the predictive power of the PGS in the HRS^47^. To facilitate comparable polygenic scores with the HRS derived PGS^48^ we omitted the MHC region on chromosome 6 from the construction of the PGS and we excluded respondents that did not meet their quality control filters. We used the HRS supplied first 10 principal components for both ego and the partner to account for population stratification^48, 49^.

### Analyses

We present two main analyses. First, we examine SGEs by simultaneously estimating the effect of ego’s and partners’ PGS on each phenotype using random effect regression models. The estimate of ego’s PGS shows how well the PGSs are able to predict health behavior in this sample. The cross-effect of the partner’s PGS on ego’s behavior net of ego’s PGS estimates the SGE. The model controls for genetic selection by including ego’s PGS, which rules out genetic selection for genetic predispositions associated with the respective health behaviors. Note that the analyses control for PCs of both ego and the partner, which provides a broad control for genetic similarity between partners. Individuals can occur as both ego and partner in the data, so each couple can occur twice (directed dyads). We adjust for repeated observations within couples over time by estimating a random intercept for dyads (directed). We use robust standard errors clustered on the household level based on sandwich estimators^50^ because husband and wife can both be ego and the partner and some individuals have had more than 1 partner in the course of the study. We furthermore control for ego’s sex (which also captures the sex of the partner because we only include heterosexual couples), both ego’s and partner’s age (linear and squared), sex-by-age interactions, and year of observation dummies and relationship length to capture secular trends and main demographic differences in health behavior.

Second, we repeat the previous analyses conditioning on initial levels of BMI/DPW/CPD for ego and partner in a couple. This second approach offers a conservative strategy to control for selection processes not controlled for using ego’s PGS and the PCs. In addition, whereas the first analysis looks at the overall level of health behavior the second analysis yields insight into the relative role of one’s own and the partner’s genes for *change* in health behavior in late adulthood. All models were also estimated for husbands and wives separately to investigate whether results were mainly driven by husbands influencing wives or vice versa. And we investigated whether there were significant interactions between ego’s and partner’s PGS, to examine social epistatic effects (i.e., genetic interaction).

To further control for selection we carried out additional analyses that adjust for well-known patterns in assortative mating (see SI for full results). We include educational level and interactions between education and sex, and mean parental education for both partners. There is ample evidence that education plays an important role in partner selection^27, 51^, and there are large educational gradients in health behaviors^38, 52^. We further include controls for height of both partners and interactions with sex, because it is also well known that height plays a role in partner selection^28, 53^. We furthermore control for region of birth of both partners to take geographic proximity into account. See SI for details.

We conducted sensitivity analyses to evaluate whether findings are affected by outliers in BMI (<18 or >45), heavy drinkers (>3 DPW), or heavy smokers (>30 CPD). In addition, we also carried out analyses excluding non-drinkers and non-smokers. Results for these analyses were not substantially different from the main analyses (see Figure S4-S5 in the SI for details).

## Acknowledgments

This research benefitted from GWAS results made publicly available by the GIANT consortium (for BMI) and the GSCAN consortium (for DPW and CPD). It was conducted by using the Health and Retirement Study genetic data (dbGAP release 2, accession number phs000428.v2.p2), which is sponsored by the National Institute on Aging (grant numbers U01AG009740, RC2AG036495, and RC4AG039029) and was conducted by the University of Michigan.

