## Supplemental Information for "*Partners in health?* Investigating social genetic effects for married and cohabiting couples"

Contents

1. Polygenic scores and phenotypes
2. Phenotypic and genotypic partner similarity, Figure S1.
3. Descriptive statistics, Table S1.
4. Coefficient estimates underlying Figure 1 and 2, Table S2-S5.
5. Social epistatic effects, Figure S2-S3.
6. Sensitivity checks, Figure S4-S5.

### 1. Polygenic scores and phenotypes

The polygenic score (PGS) for BMI (Body Mass Index) based on the most recent GWAS[1] (Genome Wide Association Study), which meta-analyzed the Locke et al. GWAS[2] and a GWAS of the UK Biobank, accounts for 13.8% of the variation in BMI using SNPs at p<.0001 in the HRS[1]. This is more than half of the proportion that is believed to be due to additive genetic factors (SNP heritability=22.4%)[1]. The PGS for smoking (cigarettes per day, CPD) and drinking behavior (drinks per week, DPW) based on the largest GWAS using data for 1.2 million people account for about ~2% of the variation in drinking (SNP heritability=4.2%) and ~4% for smoking behavior (SNP heritability=8.0%)[3].

Using our analytical sample, the PGS showed clear associations with their respective outcomes. An increase of 1 S.D. in the PGS for BMI was associated with a 1.5 point increase in BMI, and increases of 1 S.D. in the PGS for DPW and CPD were associated with 1.2 more drinks per week and 2.3 more cigarettes per day respectively.

### 2. Phenotypic and genotypic partner similarity, Figure S1

Figure S1 presents partner similarity on the phenotypic and genetic level. As reported in the main body of the paper, we find strong similarity at the phenotypic level, but weak similarity on the genetic level for all three outcomes: body mass index (BMI), alcoholic drinks per week (DPW), and cigarettes per day (CPD).

Figure S1. Phenotypic and genotypic partner similarity for BMI (weight/height$^2$), DPW (ln(drinks/week)), and CPD (ln(cigarettes/day)).

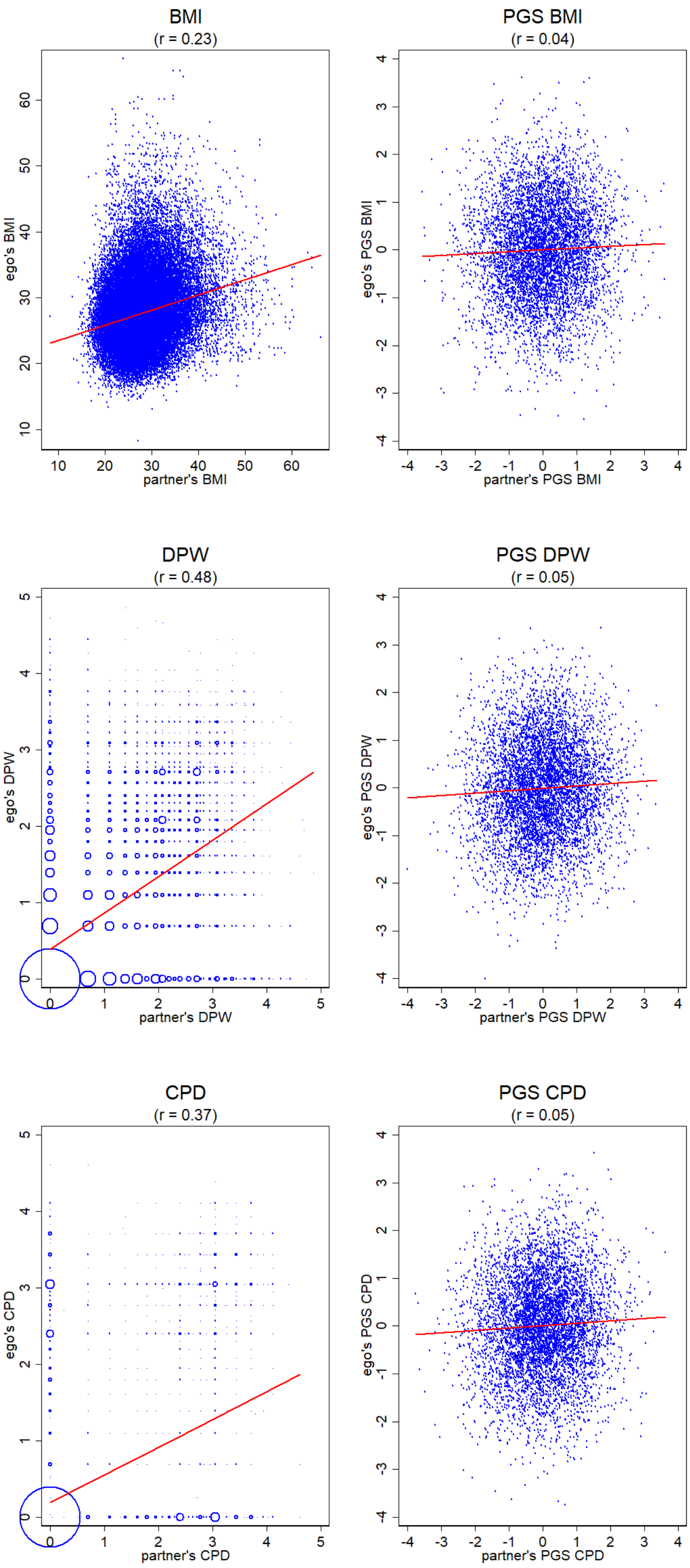


For the phenotypic scatterplots / correlations all observations were used, for the genotypic scatterplots / correlations one observation per individual. The size of the circles for DPW and CPD scale with the number of observations at each point.

#### 3. Descriptive statistics, Table S1.

The descriptive statistics are presented in Table 1. As is evident the HRS is a sample of older Americans, the average age is 65. Most relationships are long-lasting, the average duration is 34 years. The average BMI is relatively high (>25 is commonly considered overweight), and both drinking and smoking are common (respondents consume on average 3 drinks per week and 2 cigarettes per day). Comparing husbands to wives, we see that on average husbands are older (~3 years), heavier (~1 BMI point), drink more (~2 drinks more per week), but do not substantially smoke more (~⅕ cigarettes more per day).

Table S1. Descriptive statistics (means (s.d.)).

| | BMI | | | DPW | | | CPD | | |
|---|---|---|---|---|---|---|---|---|---|
| | pooled | husbands | wives | pooled | husbands | wives | pooled | husbands | wives |
| BMI (weight/height$^2$) | 27.58 | 28.03 | 27.12 | | | | | | |
| | (5.15) | (4.53) | (5.68) | | | | | | |
| drinks/week | | | | 3.00 | 3.96 | 2.04 | | | |
| | | | | (6.18) | (7.48) | (4.31) | | | |
| DPW | | | | .75 | .90 | .59 | | | |
| (ln(drinks+1/week)) | | | | (1.01) | (1.10) | (.89) | | | |
| cigarettes/day | | | | | | | 1.91 | 2.02 | 1.80 |
| | | | | | | | (6.60) | (7.09) | (6.08) |
| CPD | | | | | | | .29 | .29 | .30 |
| (ln(cigarettes+1/day)) | | | | | | | (.88) | (.89) | (.86) |
| initial outcome$_{ego, t=1}$[a] | 27.05 | 27.72 | 26.37 | 3.03 | 4.16 | 1.90 | 3.13 | 3.51 | 2.74 |
| | (4.74) | (4.09) | (5.23) | (6.04) | (7.37) | (3.99) | (8.75) | (9.67) | (7.71) |
| initial outcome$_{partner, t=1}$[a] | 27.07 | 26.42 | 27.72 | 3.04 | 1.91 | 4.17 | 3.11 | 2.72 | 3.50 |
| | (4.77) | (5.28) | (4.09) | (6.03) | (4.02) | (7.35) | (8.75) | (7.68) | (9.68) |
| age$_{ego}$ | 65.01 | 66.64 | 63.35 | 65.94 | 67.57 | 64.30 | 65.00 | 66.64 | 63.35 |
| | (9.93) | (9.61) | (9.98) | (9.76) | (9.46) | (9.77) | (9.93) | (9.61) | (9.98) |
| age$_{partner}$ | 64.98 | 63.34 | 66.64 | 65.93 | 64.32 | 67.54 | 64.99 | 63.35 | 66.64 |
| | (9.95) | (9.99) | (9.64) | (9.77) | (9.79) | (9.48) | (9.95) | (9.99) | (9.63) |
| male$_{ego}$ | .50 | | | .50 | | | .50 | | |
| | (.50) | | | (.50) | | | (.50) | | |
| PGS BMI$_{ego}$ | -.00 | .01 | -.01 | | | | | | |
| | (1.00) | (1.00) | (1.00) | | | | | | |
| PGS BMI$_{partner}$ | .00 | -.00 | .00 | | | | | | |
| | (1.00) | (.99) | (1.01) | | | | | | |
| PGS DPW$_{ego}$ | | | | -.00 | -.03 | .03 | | | |
| | | | | (1.00) | (1.01) | (.99) | | | |
| PGS DPW$_{partner}$ | | | | .00 | .03 | -.03 | | | |
| | | | | (1.00) | (.99) | (1.01) | | | |
| PGS CPD$_{ego}$ | | | | | | | -.00 | -.06 | .06 |
| | | | | | | | (1.00) | (1.01) | (.99) |
| PGS CPD$_{partner}$ | | | | | | | -.00 | .06 | -.06 |
| | | | | | | | (1.00) | (.99) | (1.01) |
| year of observation | 2004.75 | 2004.75 | 2004.75 | 2006.0 | 2006.0 | 2006.0 | 2004.8 | 2004.8 | 2004.8 |
| | (6.80) | (6.80) | (6.81) | (5.75) | (5.76) | (5.74) | (6.80) | (6.79) | (6.80) |
| relationship length | 34.50 | 34.54 | 34.46 | 35.23 | 35.26 | 35.19 | 34.50 | 34.53 | 34.47 |
| (years) | (16.02) | (15.99) | (16.04) | (16.25) | (16.22) | (16.28) | (16.01) | (16.00) | (16.03) |
| *N* observations | 52272 | 26365 | 25907 | 46734 | 23404 | 23330 | 52786 | 26394 | 26392 |
| *N* individuals | 6084 | 3043 | 3041 | 6031 | 3015 | 3016 | 6087 | 3044 | 3043 |
| *N* households | 2998 | 2997 | 2995 | 2987 | 2985 | 2986 | 2998 | 2998 | 2997 |

[a] reduced sample size for the initial outcome variables, *N*=45.626 for BMI, *N*=40.309 for DPW, *N*=47.051 for CPD.

### 4. Coefficient estimates underlying Figure 1 and 2, Table S2-S5.

Table S2. Random effects regression of BMI, DPW, CPD (t) (std.) on ego's and the partner's PGS. Standard errors adjusted for clustering in individuals and households.

| | (1) $PGS_{partner}$ - controls | (2) $PGS_{ego}$ - controls | (3) simultaneous PGS | (4) Assortative mating | (5) simultaneous PGS + interaction | (6) Assortative mating + interaction |
|---|---|---|---|---|---|---|
| BMI ($N$=52,272) | | | | | | |
| PGS $BMI_{partner}$ (std.) | .065*** | | .053*** | .044*** | .053*** | .045*** |
| | (.014) | | (.013) | (.013) | (.013) | (.013) |
| PGS $BMI_{ego}$ (std.) | | .334*** | .331*** | .319*** | .332*** | .319*** |
| | | (.013) | (.013) | (.013) | (.013) | (.013) |
| PGS $BMI_{ego}$ (std.) * | | | | | .025* | .024* |
| PGS $BMI_{partner}$ (std.) | | | | | (.012) | (.012) |
| DPW ($N$=46,734) | | | | | | |
| PGS $DPW_{partner}$ (std.) | .108*** | | .097*** | .093*** | .098*** | .094*** |
| | (.012) | | (.011) | (.011) | (.011) | (.011) |
| PGS $DPW_{ego}$ (std.) | | .210*** | .206*** | .200*** | .207*** | .200*** |
| | | (.012) | (.011) | (.011) | (.011) | (.011) |
| PGS $DPW_{ego}$ (std.) * | | | | | .022 | .018 |
| PGS $DPW_{partner}$ (std.) | | | | | (.013) | (.013) |
| CPD ($N$=52,786) | | | | | | |
| PGS $CPD_{partner}$ (std.) | .059*** | | .055*** | .034** | .055*** | .034** |
| | (.013) | | (.012) | (.012) | (.012) | (.012) |
| PGS $CPD_{ego}$ (std.) | | .079*** | .076*** | .055*** | .076*** | .055*** |
| | | (.012) | (.012) | (.012) | (.012) | (.012) |
| PGS $CPD_{ego}$ (std.) * | | | | | .020 | .022 |
| PGS $CPD_{partner}$ (std.) | | | | | (.014) | (.014) |
| controls | Yes | Yes | Yes | Yes | Yes | Yes |
| year dummies | Yes | Yes | Yes | Yes | Yes | Yes |
| $PC's_{ego}$ | No | Yes | Yes | Yes | Yes | Yes |
| $PC's_{partner}$ | Yes | No | Yes | Yes | Yes | Yes |
| assortative mating controls | No | No | No | Yes | No | Yes |

Control variables were sex, age, age$^2$ of both partners and interactions with age, age$^2$, year of observation dummies, and relationship duration. PC's were the first 10 principal components. Assortative mating controls were education (interacted with sex), height (interacted with sex), mean parental education, region of birth of both partners.

$^{\#}$ $p < .10$, $^{*}$ $p < .05$, $^{**}$ $p < .01$, $^{***}$ $p < .001$

Table S3. Sex-stratified random effects regression of BMI, DPW, CPD (t) (std.) on ego's and the partner's PGS. Standard errors adjusted for clustering in individuals and households.

| | | (1)<br>PGS$_{partner}$ + controls | (2)<br>PGS$_{ego}$ + controls | (3)<br>simultaneous PGS | (4)<br>Assortative mating | (5)<br>simultaneous PGS + interaction | (6)<br>Assortative mating + interaction |
|---|---|---|---|---|---|---|---|
| **BMI** | **men** (*N*=26,365**)** | | | | | | |
| | PGS BMI$_{partner}$ (std.) | .074$^{***}$ | | .063$^{***}$ | .060$^{***}$ | .063$^{***}$ | .060$^{***}$ |
| | | (.016) | | (.015) | (.015) | (.015) | (.015) |
| | PGS BMI$_{ego}$ (std.) | | .297$^{***}$ | .294$^{***}$ | .291$^{***}$ | .295$^{***}$ | .291$^{***}$ |
| | | | (.017) | (.017) | (.017) | (.017) | (.017) |
| | PGS BMI$_{ego}$ (std.) * | | | | | .016 | .015 |
| | PGS BMI$_{partner}$ (std.) | | | | | (.014) | (.014) |
| | **women** (*N*=25,907**)** | | | | | | |
| | PGS BMI$_{partner}$ (std.) | .057$^{**}$ | | .043$^{*}$ | .029 | .043$^{*}$ | .029 |
| | | (.021) | | (.020) | (.020) | (.021) | (.020) |
| | PGS BMI$_{ego}$ (std.) | | .372$^{***}$ | .368$^{***}$ | .347$^{***}$ | .368$^{***}$ | .348$^{***}$ |
| | | | (.020) | (.020) | (.020) | (.020) | (.020) |
| | PGS BMI$_{ego}$ (std.) * | | | | | .033$^{\#}$ | .034$^{\#}$ |
| | PGS BMI$_{partner}$ (std.) | | | | | (.017) | (.018) |
| **DPW** | **men** (*N*=23,404) | | | | | | |
| | PGS DPW$_{partner}$ (std.) | .098$^{***}$ | | .084$^{***}$ | .078$^{***}$ | .086$^{***}$ | .080$^{***}$ |
| | | (.018) | | (.018) | (.017) | (.018) | (.017) |
| | PGS DPW$_{ego}$ (std.) | | .245$^{***}$ | .242$^{***}$ | .237$^{***}$ | .243$^{***}$ | .238$^{***}$ |
| | | | (.017) | (.017) | (.017) | (.017) | (.017) |
| | PGS DPW$_{ego}$ (std.) * | | | | | .023 | .018 |
| | PGS DPW$_{partner}$ (std.) | | | | | (.017) | (.016) |
| | **women** (*N* =23,330) | | | | | | |
| | PGS DPW$_{partner}$ (std.) | .116$^{***}$ | | .108$^{***}$ | .106$^{***}$ | .109$^{***}$ | .106$^{***}$ |
| | | (.014) | | (.014) | (.013) | (.014) | (.013) |
| | PGS DPW$_{ego}$ (std.) | | .175$^{***}$ | .170$^{***}$ | .161$^{***}$ | .171$^{***}$ | .162$^{***}$ |
| | | | (.015) | (.015) | (.014) | (.015) | (.014) |
| | PGS DPW$_{ego}$ (std.) * | | | | | .019 | .017 |
| | PGS DPW$_{partner}$ (std.) | | | | | (.014) | (.013) |
| **CPD** | **men** (*N* =26,394) | | | | | | |
| | PGS CPD$_{partner}$ (std.) | .059$^{***}$ | | .055$^{**}$ | .031$^{\#}$ | .056$^{**}$ | .032$^{\#}$ |
| | | (.017) | | (.017) | (.017) | (.017) | (.017) |
| | PGS CPD$_{ego}$ (std.) | | .089$^{***}$ | .085$^{***}$ | .069$^{***}$ | .084$^{***}$ | .068$^{***}$ |
| | | | (.017) | (.017) | (.017) | (.017) | (.017) |
| | PGS CPD $_{ego}$ (std.) * | | | | | .025 | .026$^{\#}$ |
| | PGS CPD$_{partner}$ (std.) | | | | | (.016) | (.016) |
| | **women** (*N* =26,392) | | | | | | |
| | PGS CPD$_{partner}$ (std.) | .060$^{***}$ | | .055$^{**}$ | .038$^{*}$ | .055$^{**}$ | .038$^{*}$ |
| | | (.017) | | (.017) | (.017) | (.017) | (.016) |
| | PGS CPD$_{ego}$ (std.) | | .069$^{***}$ | .066$^{***}$ | .041$^{*}$ | .066$^{***}$ | .042$^{*}$ |
| | | | (.017) | (.017) | (.016) | (.017) | (.017) |
| | PGS CPD $_{ego}$ (std.) * | | | | | .016 | .018 |
| | PGS CPD$_{partner}$ (std.) | | | | | (.017) | (.017) |
| controls | | Yes | Yes | Yes | Yes | Yes | Yes |
| year dummies | | Yes | Yes | Yes | Yes | Yes | Yes |
| PC's ego | | No | Yes | Yes | Yes | Yes | Yes |
| PC's partner | | Yes | No | Yes | Yes | Yes | Yes |
| assortative mating controls | | No | No | No | Yes | No | Yes |

Control variables were sex, age, age$^2$ of both partners and interactions with age, age$^2$, year of observation dummies, and relationship duration. PC's were the first 10 principal components. Assortative mating controls were education (interacted with sex), height (interacted with sex), mean parental education, region of birth of both partners.

$^{\#}p < .10$, $^{*}p < .05$, $^{**}p < .01$, $^{***}p < .001$

Table S4. Random effects regression of BMI, DPW, CPD (t) (std.) on ego's and the partner's PGS conditioned on first observed BMI/DPW/CPD of both ego and the partner. Standard errors adjusted for clustering in individuals and households.

| | (1) $PGS_{partner}$ - controls | (2) $PGS_{ego}$ - controls | (3) simultaneous PGS | (4) Assortative mating | (5) simultaneous PGS + interaction | (6) Assortative mating + interaction |
|---|---|---|---|---|---|---|
| **BMI** (*N*=45,626) | | | | | | |
| PGS $BMI_{partner}$ (std.) | .014* | | .012# | .011# | .012# | .011# |
| | (.006) | | (.006) | (.006) | (.006) | (.006) |
| PGS $BMI_{ego}$ (std.) | | .043*** | .042*** | .042*** | .042*** | .042*** |
| | | (.007) | (.007) | (.007) | (.007) | (.007) |
| $BMI_{ego, t=1}$ (std.) | .829*** | .818*** | .817*** | .814*** | .818*** | .814*** |
| | (.009) | (.009) | (.009) | (.009) | (.009) | (.009) |
| $BMI_{partner, t=1}$ (std.) | .005 | .009 | .006 | .003 | .006 | .003 |
| | (.006) | (.006) | (.006) | (.007) | (.006) | (.007) |
| PGS $BMI_{ego}$ (std.) * | | | | | -.005 | -.005 |
| PGS $BMI_{partner}$ (std.) | | | | | (.006) | (.006) |
| **DPW** (*N*=40,309) | | | | | | |
| PGS $DPW_{partner}$ (std.) | .023*** | | .021** | .022*** | .021** | .023*** |
| | (.007) | | (.007) | (.007) | (.007) | (.007) |
| PGS $DPW_{ego}$ (std.) | | .064*** | .064*** | .065*** | .064*** | .065*** |
| | | (.007) | (.007) | (.007) | (.007) | (.007) |
| $DPW_{ego, t=1}$ (std.) | .663*** | .651*** | .649*** | .640*** | .649*** | .640*** |
| | (.010) | (.010) | (.010) | (.010) | (.010) | (.010) |
| $DPW_{partner, t=1}$ (std.) | .110*** | .115*** | .111*** | .101*** | .111*** | .100*** |
| | (.009) | (.009) | (.009) | (.009) | (.009) | (.009) |
| PGS $DPW_{ego}$ (std.) * | | | | | .005 | .006 |
| PGS $DPW_{partner}$ (std.) | | | | | (.007) | (.007) |
| **CPD** (*N*=47,051) | | | | | | |
| PGS $CPD_{partner}$ (std.) | .013# | | .013# | .011 | .013# | .011 |
| | (.007) | | (.007) | (.007) | (.007) | (.007) |
| PGS $CPD_{ego}$ (std.) | | .007 | .007 | .005 | .007 | .005 |
| | | (.007) | (.007) | (.007) | (.007) | (.007) |
| $CPD_{ego, t=1}$ (std.) | .679*** | .679*** | .678*** | .675*** | .678*** | .675*** |
| | (.015) | (.015) | (.015) | (.015) | (.015) | (.015) |
| $CPD_{partner, t=1}$ (std.) | .052*** | .053*** | .052*** | .050*** | .052*** | .049*** |
| | (.011) | (.011) | (.011) | (.011) | (.011) | (.011) |
| PGS $CPD_{ego}$ (std.) * | | | | | .014* | .014* |
| PGS $CPD_{partner}$ (std.) | | | | | (.007) | (.007) |
| controls | Yes | Yes | Yes | Yes | Yes | Yes |
| year dummies | Yes | Yes | Yes | Yes | Yes | Yes |
| PC's$_{ego}$ | No | Yes | Yes | Yes | Yes | Yes |
| PC's$_{partner}$ | Yes | No | Yes | Yes | Yes | Yes |
| assortative mating controls | No | No | No | Yes | No | Yes |

Control variables were sex, age, $age^2$ of both partners and interactions with age, $age^2$, year of observation dummies, and relationship duration. PC's were the first 10 principal components. Assortative mating controls were education (interacted with sex), height (interacted with sex), mean parental education, region of birth of both partners.

$^{\#}p < .10$, $^{*}p < .05$, $^{**}p < .01$, $^{***}p < .001$

Table S5. Sex-stratified random effects regression of BMI, DPW, CPD (t) (std.) on ego's and the partner's PGS and conditioned on first observed BMI/DPW/CPD of both ego and the partner. Standard errors adjusted for clustering in individuals and households.

| | | (1) $PGS_{partner}$ - controls | (2) $PGS_{ego}$ - controls | (3) simultaneous PGS | (4) Assortative mating | (5) sim. PGS + interaction | (6) Ass. mating + interaction |
|---|---|---|---|---|---|---|---|
| **BMI – men** | PGS $BMI_{partner}$ (std.) | .023$^{**}$ | | .022$^{**}$ | .022$^{**}$ | .022$^{**}$ | .022$^{**}$ |
| (*N*=22,931) | | (.008) | | (.008) | (.008) | (.008) | (.008) |
| | PGS $BMI_{ego}$ (std.) | | .032$^{***}$ | .031$^{***}$ | .030$^{***}$ | .030$^{***}$ | .030$^{***}$ |
| | | | (.009) | (.009) | (.009) | (.009) | (.009) |
| | $BMI_{ego, t=1}$ (std.) | .827$^{***}$ | .815$^{***}$ | .816$^{***}$ | .816$^{***}$ | .816$^{***}$ | .816$^{***}$ |
| | | (.012) | (.013) | (.013) | (.013) | (.013) | (.013) |
| | $BMI_{partner, t=1}$ (std.) | .004 | .011$^{\#}$ | .005 | .002 | .005 | .002 |
| | | (.007) | (.006) | (.007) | (.007) | (.007) | (.007) |
| | PGS $BMI_{ego}$ (std.) * PGS $BMI_{partner}$ (std.) | | | | | -.005 | -.004 |
| | | | | | | (.007) | (.007) |
| **BMI – women** | PGS $BMI_{partner}$ (std.) | .005 | | .003 | .001 | .002 | .001 |
| (*N*=22,695) | | (.010) | | (.010) | (.010) | (.010) | (.010) |
| | PGS $BMI_{ego}$ (std.) | | .054$^{***}$ | .054$^{***}$ | .053$^{***}$ | .054$^{***}$ | .052$^{***}$ |
| | | | (.010) | (.010) | (.010) | (.010) | (.010) |
| | $BMI_{ego, t=1}$ (std.) | .832$^{***}$ | .820$^{***}$ | .819$^{***}$ | .813$^{***}$ | .819$^{***}$ | .814$^{***}$ |
| | | (.012) | (.013) | (.013) | (.013) | (.013) | (.013) |
| | $BMI_{partner, t=1}$ (std.) | .006 | .007 | .007 | .006 | .007 | .006 |
| | | (.013) | (.013) | (.013) | (.013) | (.013) | (.013) |
| | PGS $BMI_{ego}$ (std.) * PGS $BMI_{partner}$ (std.) | | | | | -.008 | -.007 |
| | | | | | | (.009) | (.009) |
| **DPW – men** | PGS $DPW_{partner}$ (std.) | .015 | | .013 | .014 | .013 | .015 |
| (*N*=20,168) | | (.011) | | (.010) | (.010) | (.011) | (.010) |
| | PGS $DPW_{ego}$ (std.) | | .072$^{***}$ | .073$^{***}$ | .074$^{***}$ | .073$^{***}$ | .074$^{***}$ |
| | | | (.011) | (.011) | (.011) | (.011) | (.011) |
| | $DPW_{ego, t=1}$ (std.) | .630$^{***}$ | .618$^{***}$ | .616$^{***}$ | .614$^{***}$ | .616$^{***}$ | .614$^{***}$ |
| | | (.014) | (.014) | (.014) | (.014) | (.014) | (.014) |
| | $DPW_{partner, t=1}$ (std.) | .179$^{***}$ | .183$^{***}$ | .179$^{***}$ | .160$^{***}$ | .179$^{***}$ | .159$^{***}$ |
| | | (.017) | (.017) | (.017) | (.017) | (.017) | (.017) |
| | PGS $DPW_{ego}$ (std.) * PGS $DPW_{partner}$ (std.) | | | | | .009 | .010 |
| | | | | | | (.010) | (.010) |
| **DPW – women** | PGS $DPW_{partner}$ (std.) | .028$^{**}$ | | .026$^{**}$ | .029$^{***}$ | .026$^{**}$ | .029$^{***}$ |
| (*N*=20,141) | | (.009) | | (.009) | (.008) | (.009) | (.009) |
| | PGS $DPW_{ego}$ (std.) | | .053$^{***}$ | .053$^{***}$ | .054$^{***}$ | .053$^{***}$ | .054$^{***}$ |
| | | | (.009) | (.009) | (.009) | (.009) | (.009) |
| | $DPW_{ego, t=1}$ | .702$^{***}$ | .692$^{***}$ | .689$^{***}$ | .669$^{***}$ | .689$^{***}$ | .669$^{***}$ |
| | | (.015) | (.015) | (.015) | (.016) | (.015) | (.016) |
| | $DPW_{partner, t=1}$ | .065$^{***}$ | .071$^{***}$ | .066$^{***}$ | .064$^{***}$ | .066$^{***}$ | .064$^{***}$ |
| | | (.010) | (.010) | (.010) | (.010) | (.010) | (.010) |
| | PGS $DPW_{ego}$ (std.) * PGS $DPW_{partner}$ (std.) | | | | | -.000 | -.000 |
| | | | | | | (.009) | (.009) |
| **CPD – men** | PGS $CPD_{partner}$ (std.) | .026$^{*}$ | | .026$^{*}$ | .023$^{*}$ | .027$^{*}$ | .023$^{*}$ |
| (*N*=23,542) | | (.010) | | (.010) | (.010) | (.010) | (.010) |
| | PGS $CPD_{ego}$ (std.) | | .015 | .014 | .015 | .014 | .014 |
| | | | (.010) | (.010) | (.010) | (.010) | (.010) |
| | $CPD_{ego, t=1}$ (std.) | .619$^{***}$ | .619$^{***}$ | .618$^{***}$ | .616$^{***}$ | .618$^{***}$ | .615$^{***}$ |
| | | (.020) | (.021) | (.020) | (.020) | (.020) | (.020) |
| | $CPD_{partner, t=1}$ (std.) | .078$^{***}$ | .078$^{***}$ | .077$^{***}$ | .075$^{***}$ | .077$^{***}$ | .075$^{***}$ |
| | | (.017) | (.017) | (.017) | (.017) | (.017) | (.017) |
| | PGS $CPD_{ego}$ (std.) * PGS $CPD_{partner}$ (std.) | | | | | .018$^{*}$ | .018$^{*}$ |
| | | | | | | (.009) | (.009) |
| **CPD – women** | PGS $CPD_{partner}$ (std.) | -.001 | | -.001 | -.001 | -.001 | -.002 |
| (*N*=23,509) | | (.009) | | (.009) | (.009) | (.009) | (.009) |
| | PGS $CPD_{ego}$ (std.) | | -.001 | -.002 | -.006 | -.001 | -.006 |
| | | | (.009) | (.009) | (.009) | (.009) | (.009) |
| | $CPD_{ego, t=1}$ (std.) | .750$^{***}$ | .751$^{***}$ | .750$^{***}$ | .746$^{***}$ | .750$^{***}$ | .746$^{***}$ |
| | | (.020) | (.020) | (.020) | (.020) | (.020) | (.020) |
| | $CPD_{partner, t=1}$ (std.) | .027$^{*}$ | .026$^{\#}$ | .027$^{\#}$ | .024$^{\#}$ | .027$^{\#}$ | .024$^{\#}$ |
| | | (.014) | (.014) | (.014) | (.014) | (.014) | (.014) |
| | PGS $CPD_{ego}$ (std.) * PGS $CPD_{partner}$ (std.) | | | | | .010 | .011 |
| | | | | | | (.009) | (.009) |
| controls | | Yes | Yes | Yes | Yes | Yes | Yes |
| year dummies | | Yes | Yes | Yes | Yes | Yes | Yes |
| PC's ego | | No | Yes | Yes | Yes | Yes | Yes |
| PC's partner | | Yes | No | Yes | Yes | Yes | Yes |
| assortative mating controls | | No | No | No | Yes | No | Yes |

Control variables were sex, age, $age^2$ of both partners and interactions with age, $age^2$, year of observation dummies, and relationship duration. PC's were the first 10 principal components. Assortative mating controls were education (interacted with sex), height (interacted with sex), mean parental education, region of birth of both partners.

$^{\#}$ $p < .10$, $^{*}$ $p < .05$, $^{**}$ $p < .01$, $^{***}$ $p < .001$

### 5. Social epistatic effects, Figure S2-S3.

Figure S2 presents social epistatic effects (SEE, interaction effects between own and partner's genes) for BMI, DPW, and CPD without conditioning on past behavior, and Figure S3 presents SEE conditioning on past behavior. As reported in the main body of the paper, we find a positive SEE for BMI when not conditioning on past behavior (but no longer significant when conditioning on past behavior) and a positive SEE for CPD when conditioning on past behavior (but not significant without conditioning on past behavior).

Figure S2. Predicted BMI/DPW/CPD by own and the partner's PGS.

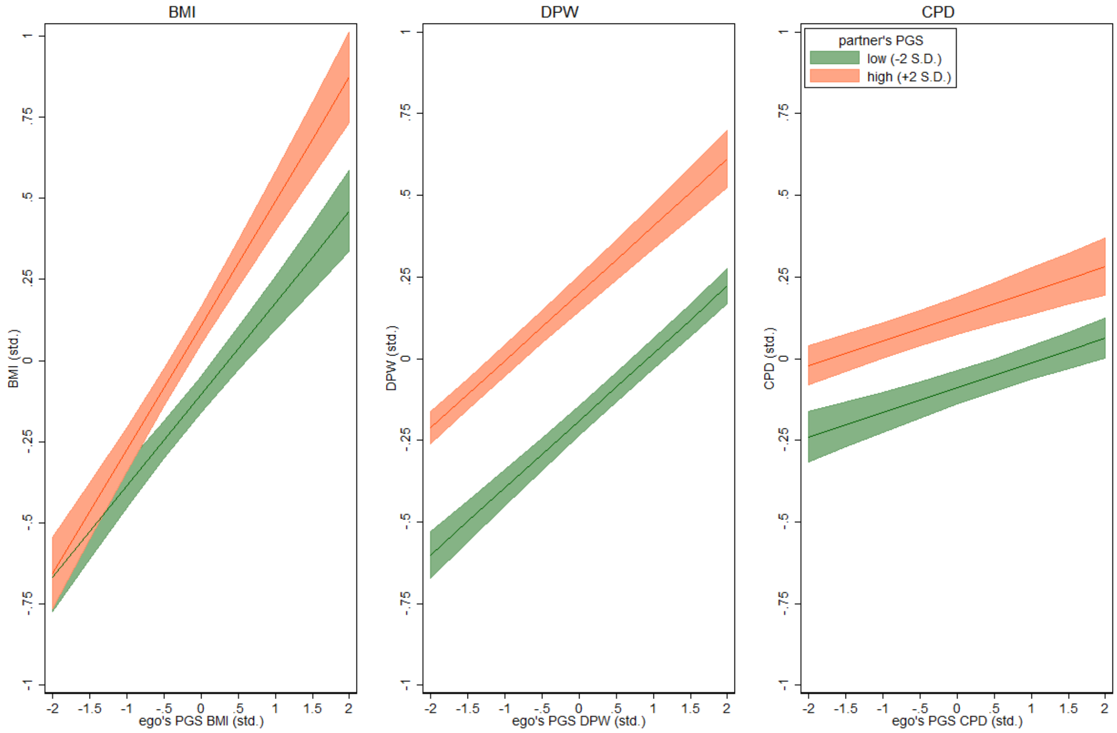


Figure S2: Marginal effects (95% CI) of ego's PGS for BMI, DPW, and CPD on associated outcomes, separating egos with partners that have low (-2 S.D.) and high (+2 S.D.) PGS. Based on a model including an interaction between own and the partner's PGS in case the social epistatic effect was statistically significant (for BMI, model 5, Table S2) or without this interaction in case the effect was not significant (for DPW and CPD, model 3, Table S2).

Figure S3. Predicted BMI/DPW/CPD by own and the partner's PGS conditioning on initial behavior of ego and partner.

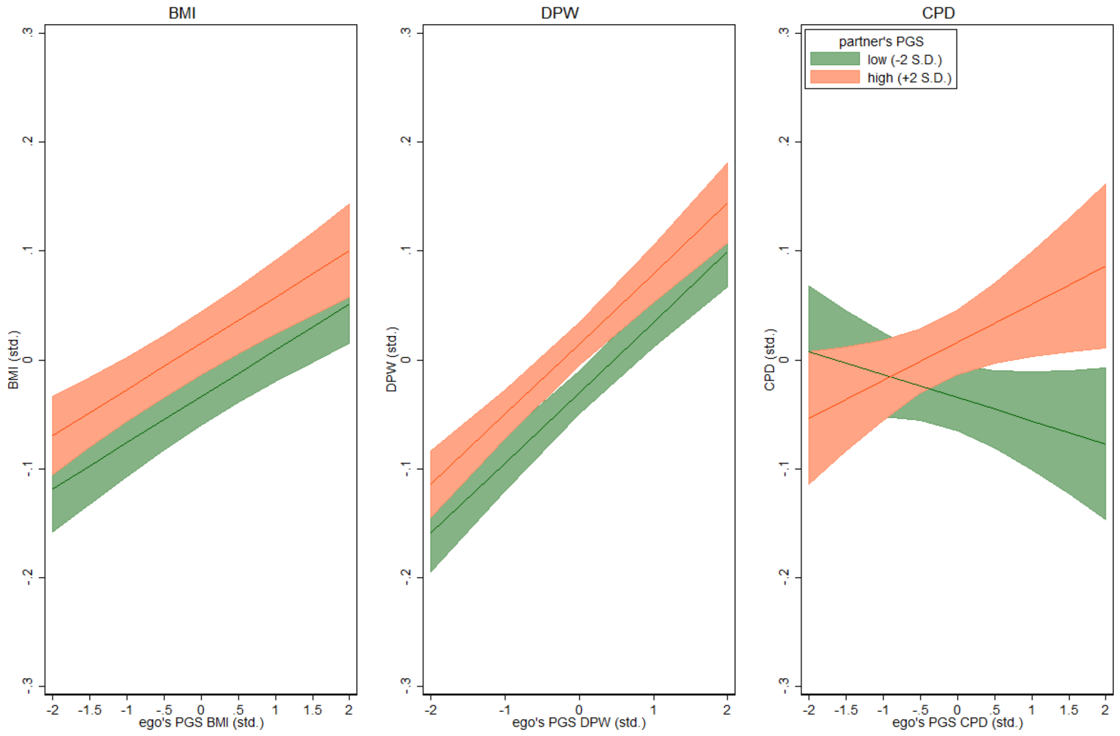


Figure S3: Marginal effects (95% CI) of ego's PGS for BMI, DPW, and CPD on associated outcomes in model controlling for initial behavior of ego and partner, separating egos with partners that have low (-2 S.D.) and high (+2 S.D.) PGS. Based on a model including an interaction between own and the partner's PGS in case the social epistatic effect was statistically significant (for CPD, model 5, Table S4) or without this interaction in case the effect was not significant (for BMI and DPW, model 3, Table S4).

### 6. Sensitivity checks, Figure S4-S5.

To assess the sensitivity of our results to outliers and sample selection criteria, we re-estimated our model that includes both ego's direct genetic effect and the partner's social genetic effect, but with the following differences:

a. Removing persons with bottom 1% and top 1% of BMI scores (BMI <18 and BMI > 45).
b. Removing persons with top 1% of CPD scores (CPD ≥ 30).
c. Removing persons with top 1-5 % of DWP scores (ln(drinks/week) ≥ 3).
d. Removing persons that never reported to have smoked.
e. Removing persons that never reported to have drunk alcoholic beverages.

Control variables were sex, age, $age^2$ of both partners and interactions with age, $age^2$, year of observation dummies, relationship duration, and the first 10 principal components for ego and partner. The results are presented in Figure S4 for analyses without conditioning on past behavior and in Figure S5 for analyses with conditioning on past behavior. The subheaders indicate which sensitivity check applies.

Figure S4. Social genetic effects of the partner, restricting sample to different subsets of respondents.

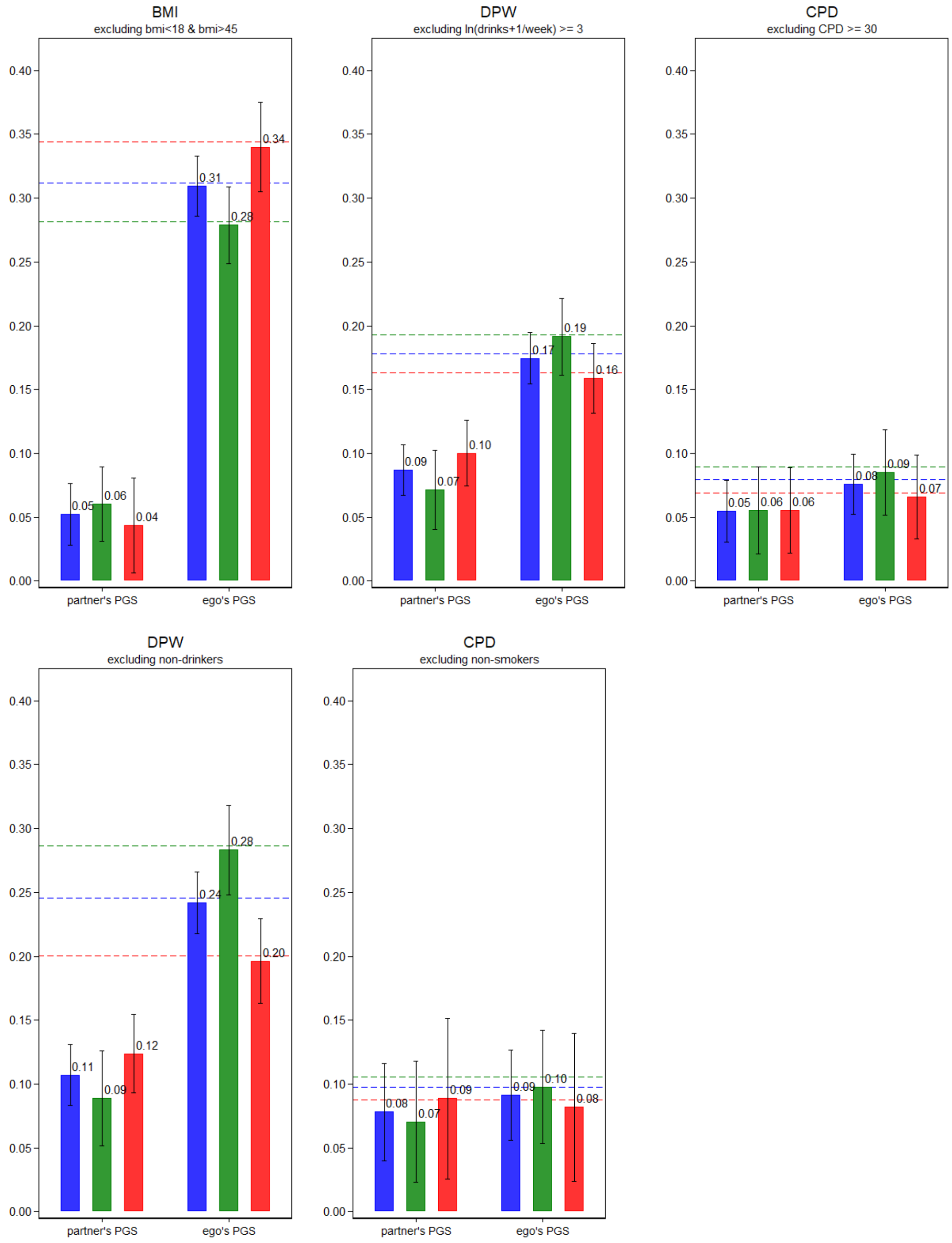


Social genetic effects of the partner. Effect of partner PGS net of one's own PGS for BMI, DPW, and CPD respectively on associated outcomes (time-varying) in a pooled model and stratified by sex with socio-demographic controls and PC's of ego and the partner. Outcomes are standardized as are the PGS. The dashed lines (blue = pooled, green = husbands, red = wives) is the baseline effect of ego's PGS on the outcome in a base model with socio-demographic controls and PC's of ego. CIs are robust to clustering within individuals and households.

Figure S5. Social genetic effects of the partner conditioning on initial behavior of ego and partner, restricting sample to different subsets of respondents.

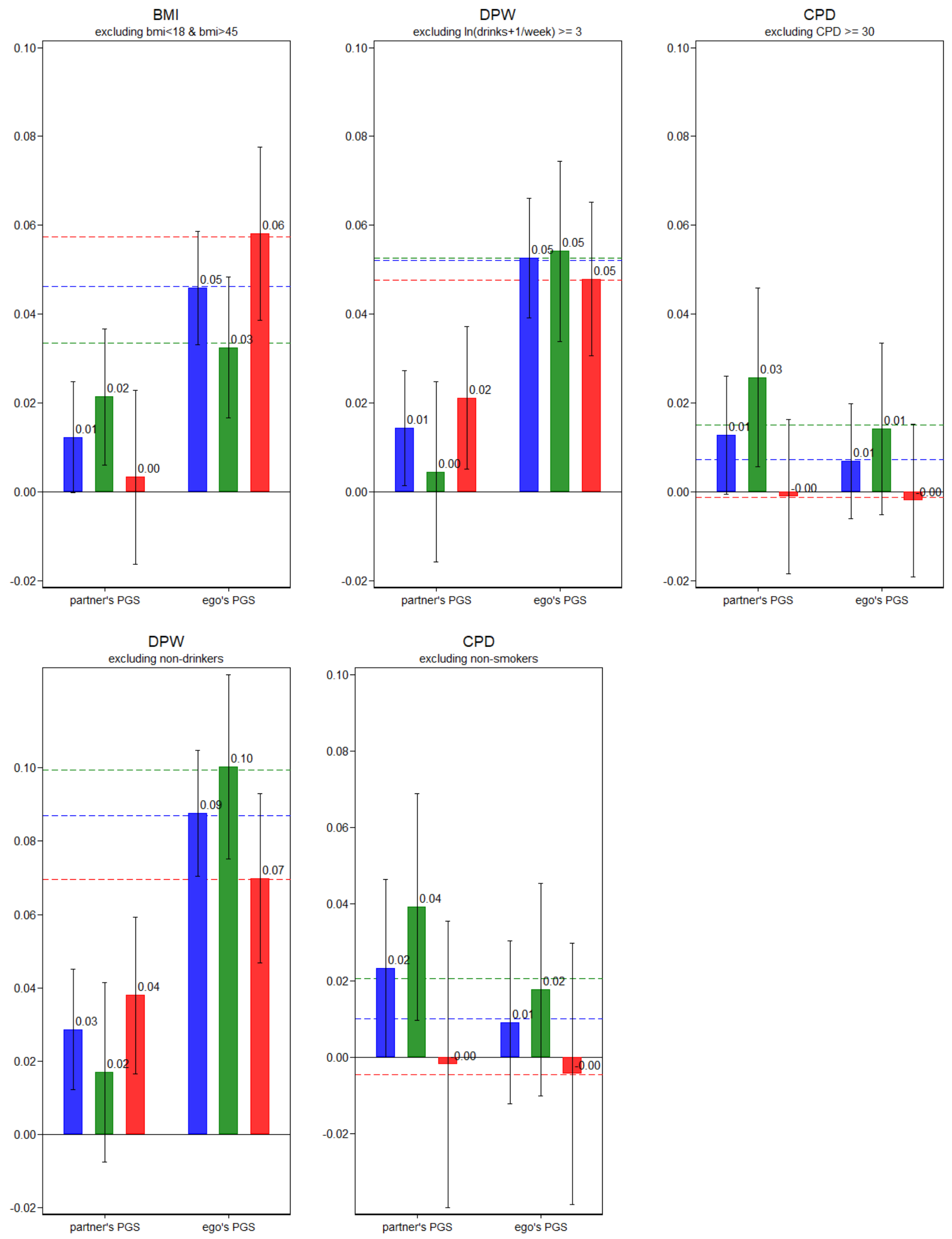


Social genetic effects of the partner conditioning on initial behavior of ego and partner. Effect of partner PGS net of one's own PGS for BMI, DPW, and CPD respectively on associated outcomes (time-varying) and net of one's own and the partner's initial level of the outcome in a pooled model and stratified by sex with socio-demographic controls and PC's of ego and the partner. Outcomes are standardized as are the PGS. The dashed lines (blue = pooled, green = husbands, red = wives) is the baseline effect of ego's PGS on the outcome in a base model with socio-demographic controls, PC's of ego and initial levels of the outcome for both ego and the partner. CIs are robust to clustering within individuals and households.
